# Wavelet Transform Coherence Under Controlled Temporal Correspondence and Social Co-Presence: Implications for fNIRS Hyperscanning

**DOI:** 10.64898/2026.09.30.755569

**Authors:** Alessandro Carollo, Frederike Soffner, Gianluca Esposito, Andrea Bizzego

## Abstract

Interpersonal neural synchrony research using functional near-infrared spectroscopy (fNIRS) frequently relies on wavelet transform coherence (WTC) to quantify temporal correspondence between cortical signals. Yet, whether WTC reliably recovers experimentally defined differences in signal correspondence remains insufficiently established. Here, we evaluated WTC using an auditory-guided finger-tapping paradigm in which matching and non-matching temporal structures were compared within participants and between dyad members during individual and joint performance. Task-evoked analyses confirmed reliable recruitment of contralateral motor and premotor regions across both contexts, whereas the ipsilateral motor cortex was significantly recruited only during joint performance and was the only region differing between sessions. WTC sensitivity was strongly modulated by social co-presence. Within participants, WTC failed to discriminate matching from non-matching temporal structures during individual performance, whereas robust matching-related increases emerged during joint performance, particularly in contralateral motor and premotor regions. Between participants, WTC showed limited and region-specific discrimination when participants were tested separately, but higher coherence for matching temporal structures emerged during joint performance, particularly in motor regions. These findings show that WTC can recover experimentally defined temporal correspondence between fNIRS signals, but that this sensitivity is not context-independent. The within-participant results further suggest that social co-presence may alter the temporal dynamics of the individual neural signals themselves. WTC should therefore be interpreted as a context-sensitive measure whose meaning depends on the functional and social processes generating the signals being compared.

## 1 Introduction

Human social interaction involves continuous coordination across individuals. During shared activities, partners may align their behavior, physiology, and neural activity over time, a phenomenon described as interpersonal synchrony [1]. Such coordination has been documented across behavioral and physiological domains, including movement, conversation, posture, and autonomic activity [2–6]. Although its expression varies across paradigms and measures [7, 8], interpersonal synchrony has been associated with several aspects of social interaction, including cooperation and social bonding [1, 9–11].

Hyperscanning has extended the study of interpersonal coordination to the neural level by enabling simultaneous recording of brain activity from multiple individuals during interaction [12, 13]. This approach has generated growing interest in interpersonal neural synchrony (INS), the temporal correspondence or coupling between neural signals recorded from different individuals [14]. Importantly, however, similar temporal dynamics across brains need not arise uniquely from interpersonal coupling. Shared sensory input, common task demands, mutual prediction, and reciprocal behavioral adaptation can all generate correspondence between neural signals [15–17]. Recent methodological arguments have further emphasized that, particularly in minimal and highly controlled paradigms, apparent INS may be strongly shaped by shared sensory, motor, and task-related processing rather than by interpersonal coupling alone [18]. Distinguishing among these sources is therefore essential for interpreting what an observed synchrony effect represents.

Functional near-infrared spectroscopy (fNIRS) is particularly suited to hyperscanning because it allows cortical hemodynamic activity to be recorded from multiple individuals under relatively unconstrained conditions [19–21]. Its portability and relative tolerance to movement have contributed to its widespread use in studies of social interaction [12, 22]. Yet, the interpretation of INS depends not only on the experimental paradigm but also on the analytical metric used to quantify correspondence between signals. Before increased synchrony can be attributed to an interpersonal process, it is therefore important to establish how the metric itself responds to experimentally defined differences in signal structure.

Several approaches have been proposed to quantify correspondence between fNIRS signals, differing in their sensitivity to temporal stability, spectral content, and lag structure [15, 22–24]. Among the most widely used is wavelet transform coherence (WTC) [11, 25, 26]. WTC characterizes the relationship between two signals in the time–frequency domain, allowing coherence to be estimated across both time and frequency [27]. Because wavelet-based approaches can accommodate changes in temporal and spectral structure, WTC is particularly attractive for naturalistic hyperscanning data [22]. However, this flexibility also makes it important to determine which properties of neural signals drive differences in WTC.

A central measurement question therefore remains: can WTC reliably distinguish signals that are expected to share temporal structure from signals that are expected to differ? Hyperscanning studies commonly evaluate INS using surrogate-dyad procedures, in which signals from individuals belonging to different dyads are artificially paired to generate a reference distribution of synchrony. Greater coherence in genuine than surrogate dyads can demonstrate that the observed correspondence is not reproduced by arbitrary pairings [28]. However, such comparisons do not establish how a synchrony metric behaves when the degree of expected temporal correspondence is experimentally manipulated. A metric may distinguish genuine from surrogate dyads while remaining sensitive to signal properties or contextual factors that are not specific to interpersonal coupling.

This issue is particularly relevant in naturalistic social interactions, where shared stimulation, common task structure, behavioral coordination, and reciprocal adaptation often occur simultaneously. Under these conditions, elevated WTC may reflect one or several sources of shared temporal structure. Controlled paradigms in which expected correspondence is defined *a priori* can therefore provide a complementary way to characterize the sensitivity of synchrony metrics and to clarify the conditions under which their output can be interpreted as evidence of INS.

The present study adopted such a bottom-up approach to evaluate WTC using fNIRS signals generated during a controlled auditory-guided finger-tapping paradigm. Rather than assuming that greater WTC necessarily reflects greater INS, we manipulated task timing to generate signal segments with different *a priori* expected levels of temporal correspondence. Participants performed 10-s finger-tapping periods in response to auditory cues, with the intervals between tapping periods systematically varied to generate distinct temporal block structures (Figure 1A–C). Finger tapping was selected because it reliably elicits hemodynamic responses in motor cortical regions [29–32], providing a predictable neural response whose temporal organization could be experimentally controlled. Blocks sharing the same temporal structure were classified as matching, whereas blocks with different temporal structures were classified as non-matching. If WTC is sensitive to experimentally imposed temporal correspondence, coherence should therefore be greater for matching than for non-matching block pairs.

**Fig. 1.**
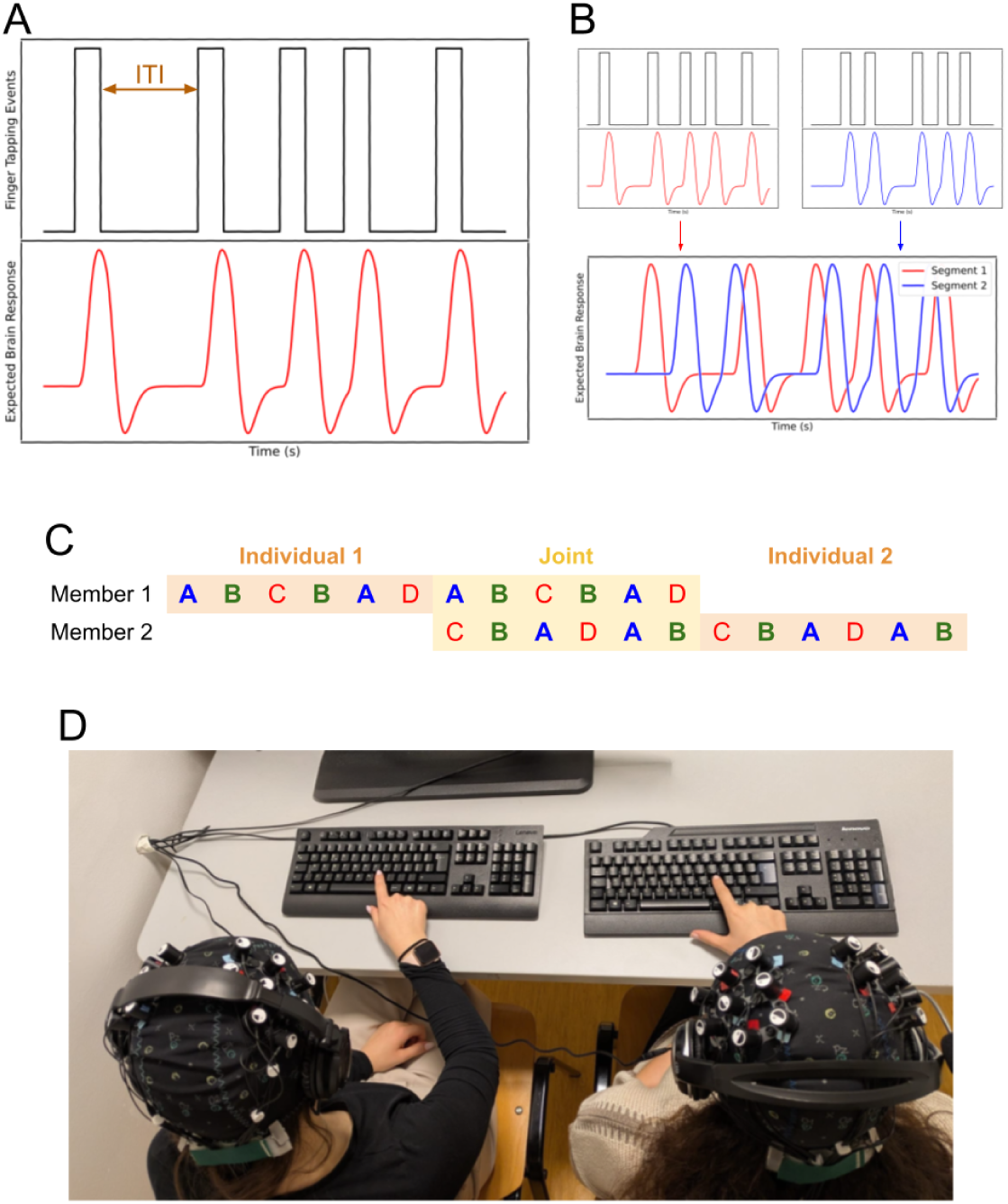
Experimental design. (A) Each block consisted of five 10-s finger-tapping periods separated by inter-tone intervals (ITIs) of variable duration, generating a predefined temporal pattern of expected hemodynamic responses. (B) Different block structures were combined to generate matching and non-matching temporal patterns. (C) Block sequences assigned to the two members of each dyad during the individual and joint sessions. (D) Experimental setting during the joint session, with both participants wearing the fNIRS cap and headphones while performing the finger-tapping task.

The design tested this measurement question across progressively more complex conditions. Within-participant comparisons provided the most controlled test by comparing signals from the same individual and minimizing inter-individual variability. Between-participant comparisons then assessed whether experimentally defined correspondence could still be recovered across members of a dyad, where individual differences in hemodynamic responses introduced additional variability. Finally, each participant performed the same assigned sequence both individually and jointly with their partner. This preserved the experimentally imposed temporal structure while introducing social co-presence, visual access to the partner’s movements, and the possibility of spontaneous mutual adaptation. Whereas the matching–non-matching manipulation provided the primary controlled test of WTC sensitivity, the individual–joint comparison assessed whether both this sensitivity and the underlying task-evoked cortical response changed when the same task was performed in the presence of a partner.

Accordingly, the study addressed two related questions. The primary aim was to determine whether WTC could discriminate between fNIRS signals characterized by experimentally defined differences in expected temporal correspondence, both within and between participants. The secondary aim was to examine whether this discrimination, together with the underlying task-evoked cortical response, differed between individual and joint performance. By separating experimentally imposed temporal correspondence from interpersonal context, this design provides a controlled test of what WTC captures and of the conditions under which WTC-based correspondence can be interpreted as evidence of INS.

## 2 Methods

The study was designed to evaluate whether WTC is sensitive to experimentally defined differences in the expected temporal correspondence between fNIRS signals and to examine how this sensitivity, together with task-evoked cortical responses, varies across individual and joint performance. Participants performed a controlled auditory-guided finger-tapping task with predefined temporal structures (Figure 1). WTC sensitivity was assessed by comparing signals associated with matching versus non-matching temporal patterns, first within participants and subsequently between members of each dyad. Each participant performed the same assigned sequence both individually and jointly with their dyadic partner, allowing the temporal manipulation to be examined in the absence and presence of social co-presence. Task-evoked cortical activation was additionally quantified to verify engagement of the targeted motor regions and to provide a functional reference for interpreting the WTC results.

### 2.1 Participants

The final sample consisted of 44 healthy adults (*M*_age_ = 25.0 years, *SD* = 7.36; 14 males, 30 females), forming 22 unique dyads. Dyad partners were not selected on the basis of age, gender, or pre-existing relationship status. Eligibility criteria required participants to be at least 18 years old and to report normal hearing. The study protocol was approved by the Institutional Ethics Committee of the University of Trento (Protocol 2023-045), and all procedures were conducted in accordance with the Declaration of Helsinki. All participants provided written informed consent prior to participation.

### 2.2 Experimental design

Each dyad completed three testing sessions: two individual sessions, one for each participant, and one joint session in which both members of the dyad performed the task simultaneously. Each participant received the same auditory sequence during their individual and joint sessions, thereby preserving the experimentally imposed temporal structure across social contexts.

During all sessions, participants wore an fNIRS cap and headphones through which auditory cues were delivered. Each auditory sequence lasted 17 min and consisted of six consecutive 170-s blocks. Each block contained five 10-s pure tones (440 Hz), separated by silent inter-tone intervals (ITIs; Figure 1A-B). Participants tapped continuously during each tone and remained still during the silent intervals. The duration and distribution of the ITIs differed across blocks, generating four predefined temporal structures (A, B, C, and D). ITIs were never shorter than 15 s, allowing the task-evoked hemodynamic response to evolve between successive tapping periods.

The four block types were arranged into two sequences: Sequence 1 (A–B–C–B–A–D) and Sequence 2 (C–B–A–D–A–B; Figure 1C). Within each dyad, one participant was assigned to Sequence 1 and the other to Sequence 2. Sequence assignment was maintained across the individual and joint sessions. Transitions between consecutive blocks occurred continuously and were not explicitly signaled to participants.

This design generated signal segments with experimentally defined differences in expected temporal correspondence. Blocks with the same temporal structure contained tapping periods at corresponding temporal positions and were therefore expected to elicit more similar temporal patterns of hemodynamic activity. In contrast, blocks with different temporal structures contained tapping periods at different temporal positions and were expected to elicit less similar hemodynamic patterns. These conditions formed the basis of the matching versus non-matching comparisons used to evaluate WTC.

Participants were instructed to tap continuously with their dominant hand on a designated keyboard key for the duration of each auditory tone and to remain still during silent intervals (Figure 1D). No instructions were provided regarding tapping rate or rhythm. During individual sessions, participants performed the task alone. During the joint session, dyad members sat side by side at the same table and could see each other’s hand movements, while direct eye contact was minimized.

Participants were informed that their auditory cues during the joint session could coincide or differ in timing. They were instructed not to communicate verbally or intentionally coordinate their behavior, and no explicit synchronization goal was imposed. Thus, the experimentally defined temporal structure remained determined by the auditory sequences, while any additional behavioral coordination associated with co-presence could emerge spontaneously.

### 2.3 fNIRS acquisition

Cortical hemodynamic activity was continuously recorded using a NIRSport2 continuous-wave fNIRS system (NIRx Medical Technologies LLC), operated with Aurora software at a sampling frequency of 10.17 Hz. For each participant, the optode montage comprised eight dual-wavelength LED sources (760 and 850 nm) and seven detectors, yielding 20 measurement channels. Optode stabilizers were used to maintain source–detector distances below 3 cm and promote stable optical coupling.

Optodes were positioned according to the International 10–20 system. Measurement channels were grouped into four regions of interest (ROIs), defined relative to the participant’s dominant hand: contralateral motor cortex (clMC), contralateral premotor cortex (clPMC), ipsilateral motor cortex (ilMC), and ipsilateral premotor cortex (ilPMC).

fNIRS acquisition, auditory stimulus presentation, and keyboard responses were synchronized in real time using the Lab Streaming Layer (LSL) protocol.

### 2.4 fNIRS signal processing

Data were processed using the *pyphysio* package [33]. All signals were resampled to 5 Hz and aligned to a common temporal reference. Separate processing pipelines were used for the task-evoked activation and WTC analyses because the two analyses addressed complementary aspects of the neural response.

#### 2.4.1 Task-evoked cortical activation

For the task-evoked activation analysis, raw light-intensity signals were converted to hemoglobin concentration changes [34]. Task-related responses were estimated separately for each participant and measurement channel using the robust general linear model framework proposed by [34], which combines optimized pre-whitening with robust regression. The resulting task-related *β* coefficients were retained as estimates of the magnitude of the cortical response to finger tapping and entered into the group-level statistical analyses.

#### 2.4.2 Preprocessing for coherence analysis

For the WTC analyses, raw light-intensity signals were first converted to optical density. Motion artifacts were identified and corrected using spline interpolation followed by wavelet-based artifact correction [35]. Corrected optical-density signals were subsequently converted to hemoglobin concentration changes. Principal component analysis filtering was applied to attenuate global physiological components [36]. Signals were then band-pass filtered between 0.01 and 0.20 Hz using an IIR filter and pre-whitened using an autoregressive model [37]. The AR model order was selected separately according to the Akaike Information Criterion, with a maximum allowed order of 10.

Signal quality was quantified separately for each measurement channel using the Scalp Coupling Index based on power estimation (SCI-P), following [38]. Rather than applying a fixed SCI-P threshold to exclude channels from the primary analyses, signal-quality estimates were incorporated into the statistical models as observation weights, as described below.

### 2.5 Wavelet transform coherence analysis

WTC was computed between pairs of preprocessed oxygenated hemoglobin (HbO) signals to quantify their time–frequency correspondence. Coherence was estimated over frequencies ranging from 0.01 to 0.20 Hz in 0.01-Hz increments.

The primary experimental factor for evaluating WTC was *block-pair correspondence*. Signal segments were classified as *matching* when they originated from blocks with the same experimentally defined temporal structure (e.g., A–A or B–B) and as *non-matching* when they originated from blocks with different temporal structures (e.g., A–C or B–D). Matching blocks were expected to generate more similar temporal patterns of task-evoked hemodynamic activity than non-matching blocks. Accordingly, greater WTC for matching than non-matching pairs constituted the primary criterion for evidence that WTC was sensitive to the experimentally imposed difference in temporal correspondence.

WTC was evaluated at two levels: within-participant and between-participant coherence.

#### 2.5.1 Within-participant coherence

Within-participant coherence was computed between HbO signals recorded from the same participant during different block occurrences. Matching comparisons paired repeated occurrences of the same block type (e.g., A_1_–A_2_ and B_1_–B_2_), whereas non-matching comparisons paired blocks characterized by different temporal structures (e.g., C–D). This analysis provided the most controlled test of WTC sensitivity because both signals originated from the same participant, thereby minimizing variability attributable to individual differences in hemodynamic response characteristics.

#### 2.5.2 Between-participant coherence

Between-participant coherence was computed between HbO signals from the two members of each dyad. Blocks were again classified as matching or non-matching according to their experimentally defined temporal structure. This analysis tested whether WTC remained sensitive to the temporal manipulation when the two signals originated from different individuals and therefore incorporated inter-individual variability in the hemodynamic response.

Both within- and between-participant analyses were performed separately for the individual and joint sessions. The individual-session analyses assessed signal correspondence in the absence of simultaneous interpersonal interaction, whereas the joint-session analyses assessed the same experimental manipulation when dyad members performed the task simultaneously in one another’s presence.

### 2.6 Statistical analyses

Linear mixed-effects models were used to account for the hierarchical structure of the data, including repeated observations within participants or dyads and multiple measurement channels within cortical regions.

#### 2.6.1 Task-evoked cortical activation

Task-related *β* coefficients were modeled to verify cortical engagement during finger tapping and to examine whether task-evoked activation differed between individual and joint sessions. Fixed effects were cortical cluster (clMC, clPMC, ilMC, ilPMC), session (individual, joint), and their interaction. Random intercepts were included for participant and for measurement channel nested within participant. Follow-up estimated marginal means were used to characterize task-evoked responses within each cluster and session and to compare activation between individual and joint sessions.

#### 2.6.2 Coherence analyses

WTC values were modeled separately for within-participant and between-participant analyses. The primary test concerned whether WTC differed between matching and non-matching block pairs. Fixed effects included cortical cluster, block-pair correspondence (matching, non-matching), and their interaction. Models were fitted separately for the individual and joint sessions. Random intercepts were included for participant in the within-participant analyses and dyad in the between-participant analyses, with measurement channel nested within participant or dyad, respectively.

Channel-level signal quality was incorporated through observation weights. For each pair of signals entering a WTC estimate, the lower of the two corresponding SCI-P values was taken as the relevant signal-quality estimate. The observation weight was defined as the square of this minimum SCI-P value. Consequently, observations based on lower-quality signal pairs contributed less to model estimation while remaining in the analysis.

Candidate model specifications were compared before conducting the primary inferential analyses. These included models without random effects, models containing participant or dyad as a random intercept, and models additionally including measurement channel nested within participant or dyad. For each random-effects specification, we compared the SCI-P weighting procedure with an alternative preprocessing strategy in which observations with SCI-P *<* 0.1 were excluded. Model performance was evaluated using the *performance* R package. Across analyses, models incorporating SCI-P-based observation weights together with participant- or dyad-level random intercepts and nested channel effects provided the best fit and were retained for the final analyses.

All linear mixed-effects models were fitted in *R* using the *lme4* and *lmerTest* packages, with sum-to-zero contrasts for categorical predictors. Fixed effects were evaluated using Type III analyses of variance with Satterthwaite approximations for denominator degrees of freedom. Follow-up comparisons were performed using *emmeans*. For the WTC analyses, estimated marginal means were used to compare matching and non-matching block pairs within each cortical cluster. *P*-values for multiple post-hoc comparisons were adjusted using the Benjamini–Hochberg false discovery rate procedure. Estimated marginal means, contrasts, and corresponding confidence intervals were reported.

## 3 Results

### 3.1 Task-evoked cortical responses

We first examined task-evoked cortical activation to verify that the finger-tapping manipulation elicited the expected hemodynamic responses and to characterize potential differences between the individual and joint sessions. The linear mixed-effects model revealed a significant main effect of cortical cluster (*F* (3, 579.63) = 11.46, *p* < .001) indicating that task-evoked responses differed across cortical regions. A significant main effect of session was also observed (*F* (1, 865.30) = 8.21, *p* = .004). These effects were qualified by a significant Cluster × Session interaction (*F* (3, 1006.72) = 2.79, *p* = .040; Table 1), indicating that session-related differences in activation varied across cortical regions.

**Table 1.** Type III analysis of variance for task-evoked cortical responses. SS = sum of squares; MS = mean square; df_1_ = numerator degrees of freedom; df_2_ = denominator degrees of freedom. * *p* < .050; ** *p* < .010; *** *p* < .001.

| Effect | SS | MS | $df_1$ | $df_2$ | $F$ | $p$ |
| --- | --- | --- | --- | --- | --- | --- |
| Cluster | .0134 | .0045 | 3 | 579.63 | 11.46 | <.001*** |
| Session | .0032 | .0032 | 1 | 865.30 | 8.21 | .004** |
| Cluster × Session | .0033 | .0011 | 3 | 1006.72 | 2.79 | .040* |

Estimated marginal means confirmed robust task-evoked activation in the contralateral motor regions (Table 2; Figure 2). In the clMC, *β* estimates were significantly greater than zero during both the individual (*M* = .1045, *SE* = .0216, *t*(92.3) = 4.83, *p* < .001) and joint sessions (*M* = .0938, *SE* = .0214, *t*(87.8) = 4.39, *p* < .001). Similarly, the clPMC showed significant positive responses during both the individual (*M* = .0455, *SE* = .0205, *t*(74.4) = 2.22, *p* = .015) and joint sessions (*M* = .0668, *SE* = .0204, *t*(73.5) = 3.27, *p* = .002).

**Fig. 2.**
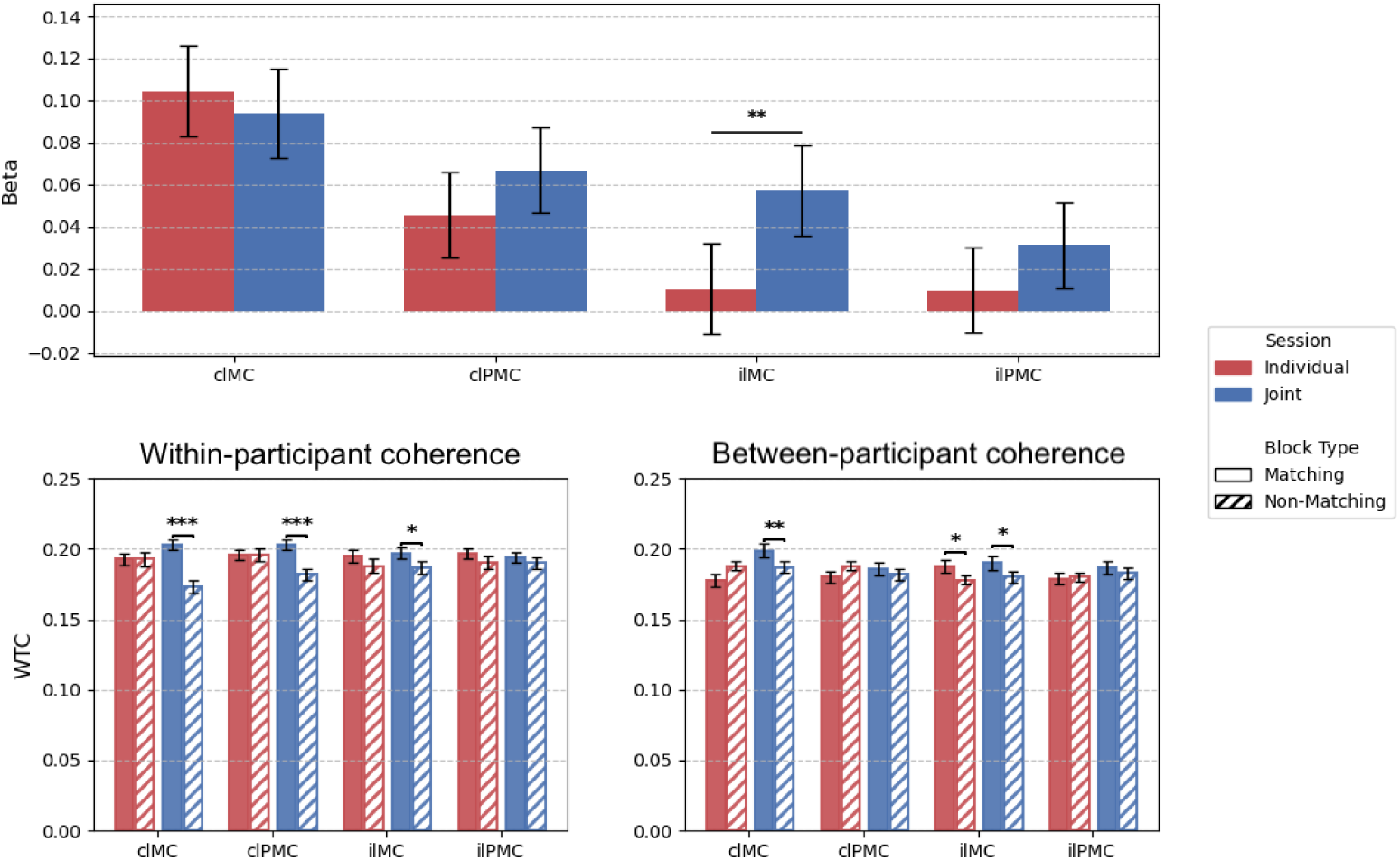
Estimated marginal means and standard errors for task-evoked cortical activation and wavelet transform coherence (WTC). Top: task-related β estimates for the individual and joint sessions. Bottom left: within-participant WTC for matching and non-matching block pairs. Bottom right: between-participant WTC for matching and non-matching block pairs. Results are shown separately for the contralateral motor cortex (clMC), contralateral premotor cortex (clPMC), ipsilateral motor cortex (ilMC), and ipsilateral premotor cortex (ilPMC). Error bars represent standard errors.

**Table 2.**
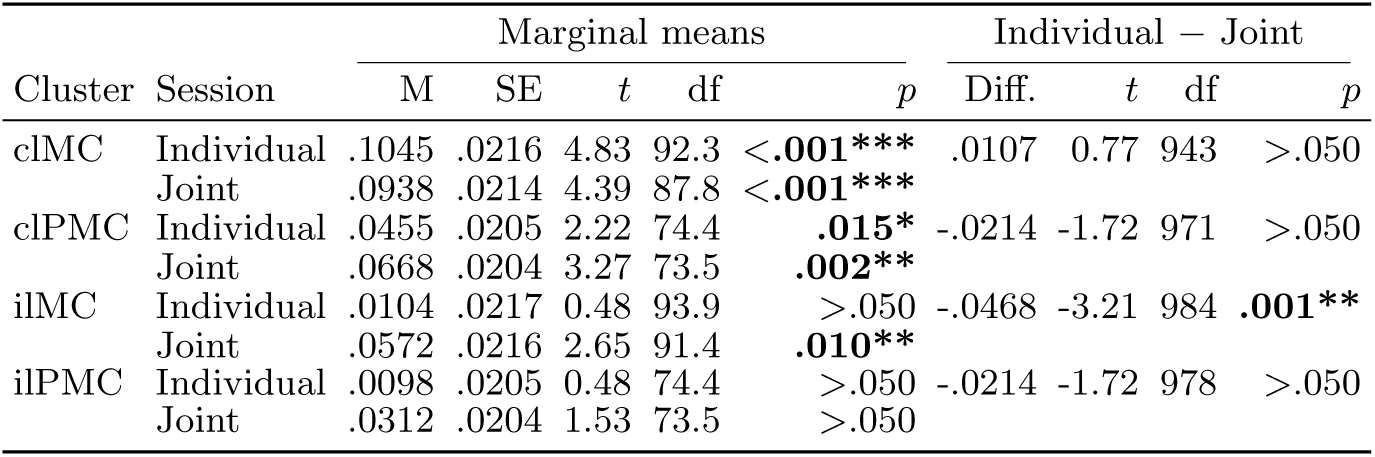
Estimated marginal means of task-evoked cortical responses and post-hoc comparisons between individual and joint sessions. clMC = contralateral motor cortex; clPMC = contralateral premotor cortex; ilMC = ipsilateral motor cortex; ilPMC = ipsilateral premotor cortex; M = estimated marginal mean; SE = standard error; Diff. = estimated difference (Individual − Joint); df = degrees of freedom. For marginal means, *t*-tests evaluate whether the estimated mean differs from zero. * *p* < .050; ** *p* < .010; *** *p* < .001.

In the ilMC, the estimated response did not differ significantly from zero during the individual session (*M* = .0104, *SE* = .0217, *t*(93.9) = 0.48, *p* > .050), whereas a significant positive response was observed during the joint session (*M* = .0572, *SE* = .0216, *t*(91.4) = 2.65, *p* = .010). The ilPMC showed no significant positive response in either the individual (*M* = .0098, *SE* = .0205, *t*(74.4) = 0.48, *p* > .050) or joint session (*M* = .0312, *SE* = .0204, *t*(73.5) = 1.53, *p* > .050).

Direct comparisons between sessions showed that activation differed significantly only in the ilMC, where *β* estimates were higher in the joint than in the individual session (Individual − Joint: Δ*M* = −.0468, *t*(984) = −3.21, *p* = .001). No significant session differences were observed in the clMC (Δ*M* = .0107, *p* > .050), clPMC (Δ*M* = −.0214, *p* > .050), or ilPMC (Δ*M* = −.0214, *p* > .050).

Together, these analyses confirmed that the finger-tapping manipulation reliably elicited the expected contralateral motor and premotor responses. They also identified a session-dependent difference in ipsilateral motor activation, providing a functional reference for the subsequent WTC analyses. The selective recruitment of the ipsilateral motor cortex during the joint session indicates that performing the same task in the presence of a partner was associated with a context-dependent modulation of cortical activity.

### 3.2 Sensitivity of WTC to experimentally defined temporal correspondence

We next examined whether WTC distinguished signal pairs characterized by experimentally defined matching versus non-matching temporal structures. This analysis was performed first within participants, providing a controlled test that minimized inter-individual variability, and subsequently between participants, where correspondence was evaluated across members of each dyad.

#### 3.2.1 Within-participant WTC

During the individual session, WTC did not reliably distinguish matching from non-matching block pairs. The Type III ANOVA showed no significant main effect of block-pair correspondence, (*F* (1, 1675.77) = 2.31, *p* > .050) and no significant main effect of cortical cluster (*F* (3, 755.87) = 0.55, *p* > .050). The Cluster × Block interaction was also not significant (*F* (3, 1757.27) = 0.70, *p* > .050; Table 3). Consistent with these results, none of the cluster-specific comparisons between matching and non-matching blocks reached statistical significance (Table 4).

**Table 3.** Type III analysis of variance for within-participant WTC in the individual and joint sessions. SS = sum of squares; MS = mean square; df_1_ = numerator degrees of freedom; df_2_ = denominator degrees of freedom. * *p* < .050; ** *p* < .010; *** *p* < .001.

| Effect | Individual |  |  |  |  |  | Joint |  |  |  |  |  |
| --- | --- | --- | --- | --- | --- | --- | --- | --- | --- | --- | --- | --- |
| | SS | MS | $df_1$ | $df_2$ | $F$ | $p$ | SS | MS | $df_1$ | $df_2$ | $F$ | $p$ |
| Cluster | .000081 | .000027 | 3 | 755.87 | 0.55 | >.050 | .000088 | .000029 | 3 | 658.41 | 0.68 | >.050 |
| Block | .000114 | .000114 | 1 | 1675.77 | 2.31 | >.050 | .002408 | .002408 | 1 | 1703.81 | 55.70 | <.001*** |
| Cluster × Block | .000104 | .000035 | 3 | 1757.27 | 0.70 | >.050 | .000897 | .000299 | 3 | 1791.49 | 6.91 | <.001*** |

**Table 4.**
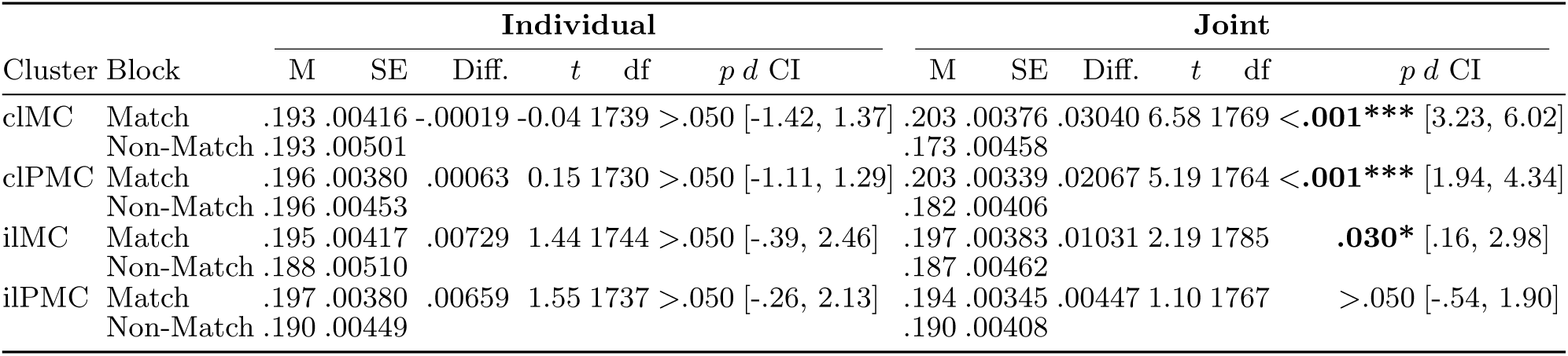
Estimated marginal means and post-hoc comparisons between matching and non-matching blocks for within-participant WTC in the individual and joint sessions. clMC = contralateral motor cortex; clPMC = contralateral premotor cortex; ilMC = ipsilateral motor cortex; ilPMC = ipsilateral premotor cortex; M = estimated maarginal mean; SE = standard error; Diff. = estimated difference (Matching − Non-Matching); df = degrees of freedom; *d* CI = lower and upper confidence limits of Cohen’s *d*. * *p* < .05; ** *p* < .01; *** *p* < .001.

A different pattern was observed during the joint session (Table 3). WTC showed a significant main effect of block-pair correspondence (*F* (1, 1703.81) = 55.70, *p* < .001), with higher coherence overall for matching than for non-matching block pairs. There was no significant main effect of cortical cluster (*F* (3, 658.41) = 0.68, *p* > .050). However, the Cluster × Block interaction was significant (*F* (3, 1791.49) = 6.91, *p* < .001), indicating that the magnitude of the matching–non-matching difference varied across cortical regions.

Follow-up comparisons showed higher WTC for matching than non-matching blocks in three of the four cortical regions. The largest difference was observed in the clMC (*M*_matching_ = .203, *M*_non-matching_ = .173, Δ*M* = .0304, *t*(1769) = 6.58, *p* < .001), followed by the clPMC (*M*_matching_ = .203, *M*_non-matching_ = .182, Δ*M* = .0207, *t*(1764) = 5.19, *p* < .001). A smaller difference was observed in the ilMC (*M*_matching_ = .197, *M*_non-matching_ = .187, Δ*M* = .0103, *t*(1785) = 2.19, *p* = .030). No significant matching–non-matching difference was observed in the ilPMC (*p* > .050).

Thus, within-participant WTC provided no evidence of discrimination between the experimentally defined temporal patterns during the individual session, whereas during the joint session it distinguished matching from non-matching blocks, particularly in contralateral motor and premotor regions.

#### 3.2.2 Between-participant WTC

We next examined whether WTC distinguished matching from non-matching temporal structures when signals originated from different members of the dyad.

During the individual session, there was no significant overall effect of block-pair correspondence (*F* (1, 1798.52) = 0.70, *p* > .050) and no significant main effect of cortical cluster (*F* (3, 388.38) = 0.65, *p* > .050). However, the Cluster × Block interaction was significant (*F* (3, 1870.98) = 3.42, *p* = .017; Table 5), indicating that the matching–non-matching difference varied across cortical regions.

**Table 5.** Type III analysis of variance for between-participant WTC in the individual and joint sessions. SS = sum of squares; MS = mean square; df_1_ = numerator degrees of freedom; df_2_ = denominator degrees of freedom. * *p* < .05; ** *p* < .01; *** *p* < .001.

| Effect | Individual |  |  |  |  |  | Joint |  |  |  |  |  |
| --- | --- | --- | --- | --- | --- | --- | --- | --- | --- | --- | --- | --- |
| | SS | MS | $df_1$ | $df_2$ | $F$ | $p$ | SS | MS | $df_1$ | $df_2$ | $F$ | $p$ |
| Cluster | .00008144 | .00002715 | 3 | 388.38 | 0.65 | >.050 | .00030150 | .00010050 | 3 | 362.40 | 2.68 | **.047\*** |
| Block | .00002924 | .00002924 | 1 | 1798.52 | 0.70 | >.050 | .00044435 | .00044435 | 1 | 1890.50 | 11.85 | **.001\*\*\*** |
| Cluster × Block | .00043106 | .00014369 | 3 | 1870.98 | 3.42 | **.017\*** | .00010409 | .00003470 | 3 | 1950.10 | 0.92 | >.050 |

Follow-up comparisons showed higher WTC for matching than non-matching blocks in the ilMC (*M*_matching_ = .188, *M*_non-matching_ = .178, Δ*M* = .0104, *t*(1857) = 2.09, *p* = .037; Table 6). No significant matching–non-matching differences were observed in the clMC, clPMC, or ilPMC (all *p* > .050). Thus, when dyad members performed the task separately, evidence that WTC distinguished the experimentally defined temporal structures was limited to a single cortical region.

**Table 6.**
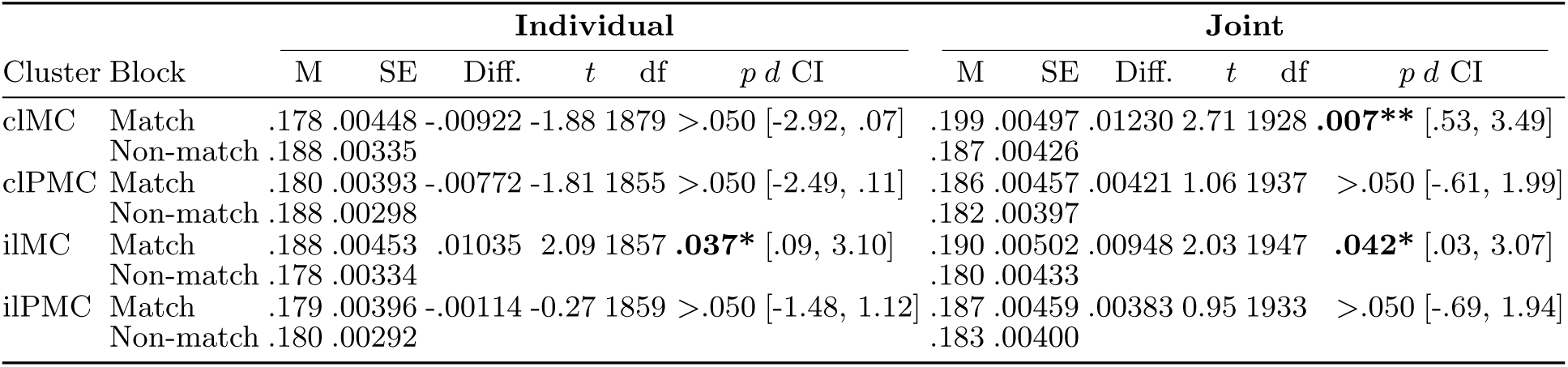
Estimated marginal means and post-hoc comparisons between matching and non-matching blocks for between-participant WTC in the individual and joint sessions. clMC = contralateral motor cortex; clPMC = contralateral premotor cortex; ilMC = ipsilateral motor cortex; ilPMC = ipsilateral premotor cortex; M = estimated marginal mean; SE = standard error; Diff. = estimated difference (Matching − Non-Matching); df = degrees of freedom; *d* CI = lower and upper confidence limits of Cohen’s *d*. * *p* < .050; ** *p* < .010; *** *p* < .001.

| Cluster Block |  | Individual |  |  |  |  |  |  | Joint |  |  |  |  |  |  |
| --- | --- | --- | --- | --- | --- | --- | --- | --- | --- | --- | --- | --- | --- | --- | --- |
| | | M | SE | Diff. | $t$ | df | $p$ | $d$ CI | M | SE | Diff. | $t$ | df | $p$ | $d$ CI |
| clMC | Match | .178 | .00448 | -.00922 | -1.88 | 1879 | >.050 | [-2.92, .07] | .199 | .00497 | .01230 | 2.71 | 1928 | **.007\*\*** | [.53, 3.49] |
|  | Non-match | .188 | .00335 |  |  |  |  |  | .187 | .00426 |  |  |  |  |  |
| clPMC | Match | .180 | .00393 | -.00772 | -1.81 | 1855 | >.050 | [-2.49, .11] | .186 | .00457 | .00421 | 1.06 | 1937 | >.050 | [-.61, 1.99] |
|  | Non-match | .188 | .00298 |  |  |  |  |  | .182 | .00397 |  |  |  |  |  |
| ilMC | Match | .188 | .00453 | .01035 | 2.09 | 1857 | **.037\*** | [.09, 3.10] | .190 | .00502 | .00948 | 2.03 | 1947 | **.042\*** | [.03, 3.07] |
|  | Non-match | .178 | .00334 |  |  |  |  |  | .180 | .00433 |  |  |  |  |  |
| ilPMC | Match | .179 | .00396 | -.00114 | -0.27 | 1859 | >.050 | [-1.48, 1.12] | .187 | .00459 | .00383 | 0.95 | 1933 | >.050 | [-.69, 1.94] |
|  | Non-match | .180 | .00292 |  |  |  |  |  | .183 | .00400 |  |  |  |  |  |

During the joint session, WTC showed a significant main effect of block-pair correspondence (*F* (1, 1890.5) = 11.85, *p* = .001; Table 5), with higher coherence overall for matching than for non-matching block pairs. A significant main effect of cortical cluster was also observed (*F* (3, 362.4) = 2.68, *p* = .047). In contrast to the individual session, the Cluster × Block interaction was not significant (*F* (3, 1950.1) = 0.92, *p* > .050), providing no evidence that the magnitude of the matching effect differed reliably across cortical regions.

Follow-up comparisons showed higher WTC for matching than non-matching blocks in the clMC (*M*_matching_ = .199, *M*_non-matching_ = .187, Δ*M* = .0123, *t*(1928) = 2.71, *p* = .007) and ilMC (*M*_matching_ = .190, *M*_non-matching_ = .180, Δ*M* = .0095, *t*(1947) = 2.03, *p* = .042). No significant differences were observed in the clPMC or ilPMC (both *p* > .050; Table 6).

Overall, the between-participant analyses provided evidence that WTC was sensitive to experimentally defined temporal correspondence, but this sensitivity depended on experimental context. When participants performed the task separately, evidence for discrimination was restricted to the ilMC and there was no overall matching effect. During joint performance, matching block pairs showed higher WTC overall, with follow-up differences observed in both contralateral and ipsilateral motor cortices.

## 4 Discussion

The present study used a controlled finger-tapping paradigm to evaluate whether WTC captures experimentally defined differences in temporal correspondence between fNIRS signals and how this sensitivity varies with social co-presence. Three main findings emerged. First, finger tapping reliably engaged contralateral motor and premotor regions across individual and joint performance, confirming the expected task-evoked response. Second, joint performance was associated with additional recruitment of the ipsilateral motor cortex, indicating that the same explicit motor task was implemented differently in the presence of a partner. Third, WTC distinguished matching from non-matching temporal structures under several conditions, but this sensitivity varied across cortical regions, within-versus between-participant comparisons, and session contexts. Matching-related differences were substantially more evident during joint than individual performance. Taken together, these findings suggest that social co-presence may influence WTC sensitivity partly by altering the neural responses from which coherence is estimated. Thus, WTC can recover experimentally defined temporal correspondence, but its expression depends on the functional and social context in which the underlying signals are generated.

The task-evoked activation results provide an important functional reference for interpreting the WTC findings. Robust activation of the contralateral motor and premotor cortices was observed during both individual and joint performance, consistent with the established involvement of these regions in voluntary finger movements [32, 39–42]. In contrast, the ipsilateral motor cortex showed significant task-evoked activation during the joint but not the individual session and was the only region showing a significant difference between the two contexts. This pattern is consistent with previous fNIRS evidence of increased recruitment of ipsilateral sensorimotor-related regions during interpersonal coordination compared with individual action execution [43], as well as reports of more bilateral cortical recruitment during interpersonal coordination [44]. Thus, although the explicit timing structure and motor requirements were unchanged, joint performance was associated with significant bilateral cortical recruitment.

Several processes may contribute to this difference. During the joint session, participants could observe their partner’s hand movements and were aware that the partner was performing a temporally related task. Joint-action contexts can modulate motor-system activity associated with observing and representing another person’s actions [45]. The joint condition may therefore have increased visuomotor processing, monitoring of external action cues, motor preparation or inhibition, or spontaneous adaptation to the partner’s behavior. Although the present design cannot distinguish among these mechanisms, the activation results indicate that social co-presence can alter the cortical implementation of an otherwise unchanged motor task.

The within-participant analysis provided the most controlled assessment of WTC sensitivity because signals were compared within the same individual, thereby minimizing variability due to stable inter-individual differences in hemodynamic responses. Given the experimentally defined temporal structure of the task, matching blocks were expected to show greater WTC than non-matching blocks in both sessions, particularly in contralateral motor regions where task-evoked activation was robust. Contrary to this expectation, WTC did not distinguish matching from non-matching temporal structures during individual performance. If experimentally imposed task timing were sufficient to produce reliable WTC discrimination, matching-related differences would be expected even when participants performed the task alone. Their absence during individual performance therefore suggests that temporal task structure alone was insufficient to produce reliable WTC discrimination. During joint performance, however, matching blocks showed significantly higher WTC in the contralateral motor and premotor cortices and, to a lesser extent, in the ipsilateral motor cortex.

This contrast across session contexts is particularly informative. The matching-related WTC differences observed during joint performance suggest that social co-presence may alter the temporal organization or consistency of the individual neural signals on which WTC is computed. This interpretation is consistent with recent methodological arguments that interbrain synchrony can partly arise from greater behavioral consistency and the resulting temporal alignment of task-related neural responses [18]. Hamilton and Konvalinka, for example, argue that more consistent behavioral timing can produce more tightly aligned neural time series, even when the observed correspondence does not necessarily reflect a uniquely interpersonal neural mechanism. In the present study, however, the experimentally imposed temporal structure was already present during individual performance, where WTC did not reliably discriminate matching from non-matching blocks. Social co-presence may therefore have increased the consistency or expression of task-related neural dynamics, making the imposed temporal correspondence more detectable by WTC.

Several processes may account for this effect. During the joint session, participants could observe their partner’s hand movements and were aware that the partner was performing a temporally related task. Visible partner behavior may have provided additional temporal information, increased attention to task timing, or promoted subtle behavioral adaptation, potentially reducing variability in motor execution and its associated hemodynamic response. Such adaptation need not have been deliberate, as participants were explicitly instructed not to coordinate their behavior. From this perspective, interpersonal context may influence synchrony estimates not only by creating relationships between participants, but also by modifying the temporal properties of the individual signals being compared.

The ipsilateral motor cortex provides a concrete example of this context dependence. During joint performance, this region showed both increased task-evoked recruitment and matching-related differences in WTC. Although these findings do not demonstrate that changes in activation caused the WTC effect, they show that the coherence difference emerged in a region whose task-related response also changed across contexts. This suggests that WTC should be interpreted in relation to the properties of the signals on which it is computed: when regional engagement or the temporal consistency of the neural response changes, the correspondence detected by WTC may also change.

The between-participant analyses provide a complementary perspective that is directly relevant to INS. When dyad members were recorded separately, their experimentally imposed task structures were preserved, but real-time co-presence, visual monitoring, and reciprocal adaptation were absent. This condition therefore provided a reference for the degree of between-participant correspondence that could arise from task structure alone. There was no overall matching effect, although a significant Cluster × Block interaction was observed and post-hoc analyses showed higher WTC for matching blocks in the ipsilateral motor cortex. Thus, measurable between-participant correspondence could emerge even in the absence of concurrent interaction, demonstrating that shared task structure can contribute to neural similarity across individuals without being sufficient to explain the full pattern observed during joint performance.

During joint performance, however, WTC was higher overall for matching than for non-matching blocks, with significant matching-related differences in both the contralateral and ipsilateral motor cortices. Particularly notable was the contralateral motor cortex, where matching-related discrimination was detected during joint but not separate performance. This pattern is consistent with the possibility that social co-presence contributes additional temporal structure beyond that imposed by the task. Visual monitoring, shared attention, or spontaneous motor adaptation could all increase the consistency of task-related neural responses and thereby make experimentally defined correspondence more detectable by WTC.

Taken together, the within- and between-participant findings support a qualified interpretation of WTC. The metric was capable of recovering experimentally defined differences in temporal correspondence, but it did not do so uniformly. Its sensitivity varied with cortical region, level of comparison, and experimental context. WTC should therefore not be treated as a context-independent or uniquely interpersonal index of neural synchrony. Rather, it quantifies time–frequency correspondence between signals whose properties are themselves shaped by the neural and behavioral context in which they are generated.

This interpretation also highlights the value of considering synchrony measures together with conventional measures of regional brain activation. The ipsilateral motor cortex provides a clear example: selective recruitment during joint performance accompanied matching-related WTC differences in the same region. Had WTC been considered in isolation, this change in the underlying cortical response would have been missed. More generally, elevated synchrony does not imply that the same neural processes are operating equivalently across conditions. Changes in regional engagement can alter the signals on which synchrony metrics are computed and, consequently, the resulting estimates of correspondence.

These considerations are particularly important for naturalistic hyperscanning, where shared sensory input, common task demands, behavioral coordination, mutual prediction, and reciprocal adaptation often coexist. Under such conditions, elevated WTC may reflect several sources of temporal correspondence simultaneously. The present findings further suggest that social context may influence synchrony estimates not only by affecting relationships between participants, but also by altering the neural responses generated within each individual. Surrogate-dyad approaches provide an important statistical reference by testing whether synchrony in genuine pairs exceeds that expected from artificial pairings, but they do not determine which features of the interaction generated that correspondence. Controlled paradigms such as the one used here address a complementary question by experimentally defining expected signal relationships while also examining how interpersonal context changes the signals on which synchrony metrics are computed. Combining controlled characterization of synchrony metrics with surrogate-based approaches may therefore provide a stronger methodological basis for distinguishing task-driven, signal-driven, and interaction-related contributions to neural synchrony in complex social settings.

### 4.1 Limitations

Some limitations should be considered. First, finger tapping represents a highly constrained motor behavior and therefore captures only a limited subset of the processes involved in natural social interaction. The extent to which the present findings generalize to more complex affective, communicative, or cooperative exchanges remains to be established. Second, the study focused on motor and premotor regions and on WTC applied to fNIRS signals; other brain systems, imaging modalities, and synchrony metrics may show different sensitivity to experimentally imposed temporal structure. Finally, the joint condition involved social co-presence and visual access to the partner’s movements but did not require explicit cooperation or synchronization. The effects observed in this condition should therefore be interpreted as modulation associated with co-presence and the possibility of mutual adaptation rather than as evidence for deliberate interpersonal coordination.

## 5 Conclusion

This study provides a controlled evaluation of WTC as a measure of temporal correspondence between fNIRS signals. By experimentally manipulating the temporal structure of a motor task, we tested whether WTC could distinguish signal pairs with different *a priori* expected levels of correspondence. WTC discriminated matching from non-matching temporal patterns under several conditions, demonstrating sensitivity to experimentally defined signal relationships. However, this sensitivity varied across cortical regions, within- versus between-participant comparisons, and experimental contexts.

These findings support a context-dependent interpretation of WTC. The metric can recover experimentally imposed differences in temporal correspondence, but its sensitivity appears to depend on the properties of the underlying neural signals and on the context in which those signals are generated. Notably, joint performance was associated with both altered regional cortical recruitment and more evident matching-related WTC differences. The within-participant findings further suggest that social co-presence may alter the temporal organization or consistency of individual neural responses themselves, thereby influencing synchrony estimates.

WTC should therefore not be interpreted as a context-independent or uniquely interpersonal index of neural synchrony. Rather, it quantifies time–frequency correspondence between signals whose structure may reflect shared task demands, regional neural recruitment, and interpersonal context. Social interaction may thus shape measured neural synchrony not only through relationships between participants, but also by modifying the individual neural signals being compared. Controlled, bottom-up paradigms such as the one adopted here can complement naturalistic hyperscanning by helping to disentangle these contributions and by strengthening the interpretation of neural synchrony in complex social interactions.

## Author contributions

Conceptualization: AC, AB; Methodology: AC, AB; Software: AB; Formal analysis: FS, AB; Investigation: FS; Data Curation: FS, AB; Writing - Original Draft: AC, FS, AB; Writing - Review & Editing: AC, AB, GE; Supervision: AB.

## Ethics declarations

The study protocol was approved by the Institutional Ethics Committee of the University of Trento (Protocol 2023-045), and all procedures were conducted in accordance with the Declaration of Helsinki. All participants provided written informed consent prior to participation.

